# Effective connectivity reveals distinctive patterns in response to others’ genuine affective experience of disgust as compared to pain

**DOI:** 10.1101/2021.09.03.458875

**Authors:** Yili Zhao, Lei Zhang, Markus Rütgen, Ronald Sladky, Claus Lamm

**Affiliations:** Social, Cognitive and Affective Neuroscience Unit, Department of Cognition, Emotion, and Methods in Psychology, Faculty of Psychology, University of Vienna, Liebiggasse 5, 1010 Vienna, Austria; Vienna Cognitive Science Hub, University of Vienna, Liebiggasse 5, 1010 Vienna, Austria

**Keywords:** empathy, emotion, recognition, insular cortex

## Abstract

Empathy is significantly influenced by the identification of others’ emotions. In a recent study, we have found increased activation in the anterior insular cortex (aIns) that could be attributed to affect sharing rather than perceptual saliency, when seeing another person genuinely experiencing pain as opposed to merely acting to be in pain. In that prior study, effective connectivity between aIns and the right supramarginal gyrus (rSMG) was revealed to track what another person really feels. In the present study, we used a similar paradigm to investigate the corresponding neural signatures in the domain of empathy for disgust - with participants seeing others genuinely sniffing unpleasant odors as compared to pretending to smell something disgusting. Consistent with the previous findings on pain, we found stronger activations in aIns associated with affect sharing for genuine disgust compared with pretended disgust. However, instead of rSMG we found engagement of the olfactory cortex. Using dynamic causal modeling (DCM), we estimated the neural dynamics of aIns and the olfactory cortex between the genuine and pretended conditions. This revealed an increased excitatory modulatory effect for genuine disgust compared to pretended disgust. For genuine disgust only, brain-to-behavior regression analyses highlighted a link between the observed modulatory effect and the perspective-taking empathic trait. Altogether, the current findings complement and expand our previous work, by showing that perceptual saliency alone does not explain responses in the insular cortex. Moreover, it reveals that different brain networks are implicated in a modality-specific way when sharing the affective experiences associated with pain vs. disgust.

## Introduction

Our affective states are remarkably affected by the perceived feelings of others. A theoretical framework of empathy proposed by Coll et al. (2017) states that identification of another’s emotion crucially contributes to the consequential sharing of feelings with that person. A recent study by our group has revealed that when individuals witness another person genuinely experiencing pain as compared to merely acting to be in pain, they attribute more painful feelings to that person and report experiencing stronger self-unpleasantness in response to the other’s genuine pain (Zhao et al., 2021b).

On the neural level, that study found increased brain activations for the genuine compared to the pretended pain in the anterior insular cortex (aIns) and the anterior mid-cingulate cortex (aMCC), i.e., a network that has been consistently associated with affective responding in studies on self- experienced pain as well as empathy for pain (Lamm et al., 2011; Rütgen et al., 2015; Jauniaux et al., 2019; Xiong et al., 2019; Feng Zhou et al., 2020; Fallon et al., 2020, for meta-analyses). One major contribution of our previous study is that we have shown aIns, a key node of this neural network, is indeed associated with affect sharing, rather than being driven by the perceptual saliency of the facial expressions of pain. Moreover, by means of dynamic causal modeling (DCM) analyses, distinctive effective connectivity of genuine pain vs. pretended pain has been found on the connection between aIns and the right supramarginal gyrus (rSMG), a region selectively related to affective self-other distinction, in the sense that it is distinct from a more posterior part centering more on the angular gyrus, the temporo-parietal junction, that is implicated in self-other distinction in the cognitive domain (Silani et al., 2013; Steinbeis et al., 2015; Hoffmann et al., 2016; Bukowski et al., 2020). This suggests that the interaction of aIns and rSMG tracks how we identify and share the actual feelings of another person, allowing an observer to engage in appropriate affect sharing rather than simply responding to salient, yet possibly non-genuine displays of pain.

What remains an open question is whether these findings are specific to pain or could be extended to other aversive experiences. Among the array of aversive experiences, the emotion of disgust partially overlaps with pain regarding its neural mechanisms (Corradi-Dell’Acqua et al., 2016). Also, disgust and pain share similarities with respect to their facial expression (Zhao et al., 2021a) and are similarly important for survival and somatic protection (Sharvit et al., 2015; Sharvit et al., 2020). Particularly, research using multi-voxel pattern analysis (MVPA) shows overlapping brain maps in aIns and aMCC not only for self-experienced but also vicarious experiences of pain and disgust, suggesting a modality-independent representation of the unpleasantness shared by self-experienced aversive affect and empathy for such affect (Corradi-Dell’Acqua et al., 2016).

The aim of the present study was, thus, to replicate and expand the findings of our previous study on pain (Zhao et al., 2021b), but targeting the emotion of disgust. Specifically, participants watched video clips either presenting a person showing a disgust expression when sniffing something unpleasant, or merely displaying a disgust expression without genuinely smelling any unpleasant odor. We expected to find that 1) on the behavioral level, genuine disgust would result in higher other-oriented disgust ratings and self-oriented unpleasantness ratings; 2) on the neural level, aIns, aMCC, and rSMG would show stronger responses to the genuine disgust, as compared to pretended disgust; and 3) distinct patterns of aIns’ effective connectivity with rSMG would be found, and explain the different empathic responses to genuine vs. pretended disgust in a similar way as for pain.

## Materials and Methods

To maximize comparability, data collection for the current study had been planned and performed together with the study focusing on pain (Zhao et al., 2021b). Thus, all procedures of both studies (i.e., creation and validation of stimuli, the pilot study, and the main fMRI experiment) were exactly conducted in the same sessions and with the identical participant sample. We decided to analyze and report them separately for reasons of reporting complexity and as the two reports have a different focus. While the details about all procedures are fully documented in (Zhao et al., 2021b), for reasons of enhancing accessibility and reproducibility, we summarize the main points relevant to the current study herein.

### Participants

Forty-eight participants participated in this study. This sample size was estimated a priori using Gpower 3.1 (Faul et al., 2007), for which a minimum sample size statistically required for this study was 34 with a medium effect size of Cohen’s d = 0.5 (α = 0.05, two-tailed, 1−β = 0.80). Three participants (only for the current study) were excluded because of excessive head motion (> 15% scans with the frame-wise displacement over 0.5 mm in one session; same criteria as the pain study). Data of the remaining 45 participants (21 females; age: Mean = 26.76 years, S.D. = 4.58) were entered into analyses. Participants had normal or corrected to normal vision and were pre-screened by an MRI safety-check questionnaire to assure no presence or history of neurologic, psychiatric, or major medical disorders. All participants reported being right-handed and signed the informed consent. The study was approved by the ethics committee of the Medical University of Vienna and was conducted in accordance with the latest revision of the Declaration of Helsinki (2013).

### Manipulation of facial expressions

We created the stimuli for disgust with the same demonstrators who had also performed for the pain stimuli. In strict analogy to the stimuli we created for pain, the stimuli we created for this study consisted of video clips showing demonstrators ostensibly in four different situations: 1) Genuine disgust: the demonstrator sniffed dog feces in an *opened* bottle with a picture depicting dog feces on it; the demonstrator’s facial expression changed from neutral to strongly disgusted. 2) Genuine no disgust: the demonstrator sniffed cotton balls in an opened bottle with a picture depicting cotton balls on it; the demonstrator’s facial expression maintained neutral. 3) Pretended disgust: the demonstrator sniffed dog feces in a *closed* bottle (covered by a cap) with a picture depicting dog feces on it; the demonstrator’s facial expression changed from neutral to strongly disgusted. 4) Pretended no disgust: the demonstrator sniffed cotton balls in an opened bottle with a picture depicting cotton balls on it; the demonstrator’s facial expression maintained neutral. In fact, the genuine no disgust condition and pretended no disgust condition contained identical videos. We randomly labeled 50% of neutral videos as “genuine no disgust” when the same demonstrators appeared in the genuine disgust condition, and treated the other 50% neutral videos as “pretended no disgust” when the same demonstrators presented in the pretended disgust condition.

Twenty demonstrators (10 females), with experience in acting, were recruited for creating the stimuli of the current study. Each demonstrator signed the agreement of using their videos clips and static images for scientific purposes. An experimenter who stood on the right side of the demonstrators, of whom only the right hand holding the bottle could be seen, moved the bottle from the demonstrator’s right side and stopped it just below the demonstrator’s nose. Unbeknownst to the participants, all disgusted expressions were acted and the so-called “dog feces” were actually an odor-neutral object that resembled dog feces. As soon as the bottle was close enough to the demonstrator’s nose (just below the right nostril), the demonstrator started to make a disgusted facial expression along with a slightly avoidant movement of their head, as naturally and vividly as possible. In the neutral control conditions, demonstrators maintained a neutral facial expression during the whole process of the bottle movement. Note that, the reason for presenting the pictures and supposed content of dog feces in both disgust conditions was because we deemed it essential to match the conditions in terms of the presence and visibility of an aversive disgusting object approaching the other person’s face. Otherwise, any difference between conditions could be confounded by responses of participants to the presence vs. absence of a disgusting object and its explicit photographic display. Note that the pain condition of our previous work also followed this logic, with a needle covered by a plastic cap approaching the cheek.

### Stimulus validation and pilot study

To validate the stimuli, 110 participants (59 females; age: Mean = 29.32 years, S.D. =10.17) were recruited and asked to rate a total of 120 video clips of 2 s duration of the two conditions (60 of each condition) showing disgusted facial expressions (i.e., the genuine and pretended disgust conditions). The main aim of the validation study was to identify a set of demonstrators that expressed disgust with comparable intensity and quality, and whose expressions of disgust in the genuine and pretended conditions were comparable. After each video clip, participants rated three questions on a visual analog scale with 9 tick-marks and the two end-points marked as “almost not at all” to “unbearable”: 1) How much disgust did the person *express* on his/her face? 2) How much disgust did the person *actually feel*? 3) How unpleasant *did you feel* to watch the person in this situation? These questions were presented in a pseudo-randomized order. Moreover, we set eight catch trials to test whether participants maintained attention to the stimuli, in which participants were required to correctly choose the demonstrator they had seen in the last video, from two static images showing either the correct demonstrator’s or a distractor’s neutral facial expression side by side.

Data collection was performed with the online survey platform SoSci Survey (https://www.soscisurvey.de), and participants got access to the survey through a participation invite published on Amazon Mechanical Turk (https://www.mturk.com/). Survey data of 62 out of 110 participants (34 females; age: Mean = 28.71 years, S.D. =10.11) were entered into the analysis (inclusion criteria: false rate for the test questions < 2/8, survey duration > 20 min and < 150 min, and the maximum number of continuous identical ratings < 5). According to the validation analysis, videos of 6 demonstrators (3 females) were excluded for which participants showed a significant difference in perceived disgust expressions in others between genuine disgust and pretended disgust. As a result of this validation, videos of 14 demonstrators (7 females) were selected for the subsequent pilot study.

In the pilot study, a separate group (N =47, 24 females; age: Mean = 26.28 years, S.D. = 8.80) were recruited for a behavioral experiment in the behavioral laboratory. All conditions including the neutral conditions described above were presented to the participants to test the feasibility of the procedures that we intended to use in the following fMRI experiment. Participants were explicitly instructed that they would watch some persons’ genuine expressions of disgust in some blocks, while in other blocks, they would see some other persons acting out disgust expressions (recall that in reality, all demonstrators had been actors). All demonstrators showed neutral expressions as well. The three questions mentioned above were required to be rated. According to the video screening, we excluded videos of two demonstrators (1 female) for whom participants showed a large difference in ratings of *expression* of disgust between pretended vs. genuine conditions. Three separate repeated-measures ANOVAs were respectively performed for the three rating questions regarding the remaining videos. For the disgusted expressions in others, the main effect of genuineness (genuine vs. pretended) was not significant and was low in effect size (*F* _genuineness_ (1, 46) = 0.867, *p* = 0.357, η^2^ = 0.018), but it was significant and showed high effect size for the disgusted feelings in others (*F* _genuineness_ (1, 46) = 207.225, *p* < 0.001, η^2^ = 0.818) as well as for the unpleasantness in self (*F* _*genuineness*_ (1, 46) =21.360, *p* < 0.001, η^2^ = 0.317). The main effects of disgust (disgust vs. no disgust) for all three ratings were significant with high effect size (the smallest effect size was for the rating of unpleasantness in self, *F* _disgust_ (1, 46) = 44.489, *p* < 0.001, η^2^ = 0.492). The findings of our pilot study for the domain of disgust were thus very much in line with the findings of the same pilot study for the domain of pain (see Zhao et al., 2021b). Finally, video clips of 12 demonstrators (6 females) were determined for the main fMRI experiment.

### Experimental design and procedures of the fMRI study

The experimental design and procedures are sketched in Figure 1A and 1B. The fMRI experiment was performed in two runs, and each run consisted of two blocks showing genuine disgust and two blocks showing pretended disgust. The order of genuine (“G”) blocks and pretended (“P”) blocks was pseudo-randomized across participants; specifically, the block order was “G G P G P G P P” for one half of the participants and “P P G P G P G G” for the other half. In each block, participants watched nine video clips containing both disgusted and neutral videos. The order of disgusted videos and neutral videos was pseudo-randomized. Additionally, the order of disgust sessions and pain sessions (with the latter being reported in Zhao et al., 2021b), was counterbalanced across participants.

**Figure 1.**
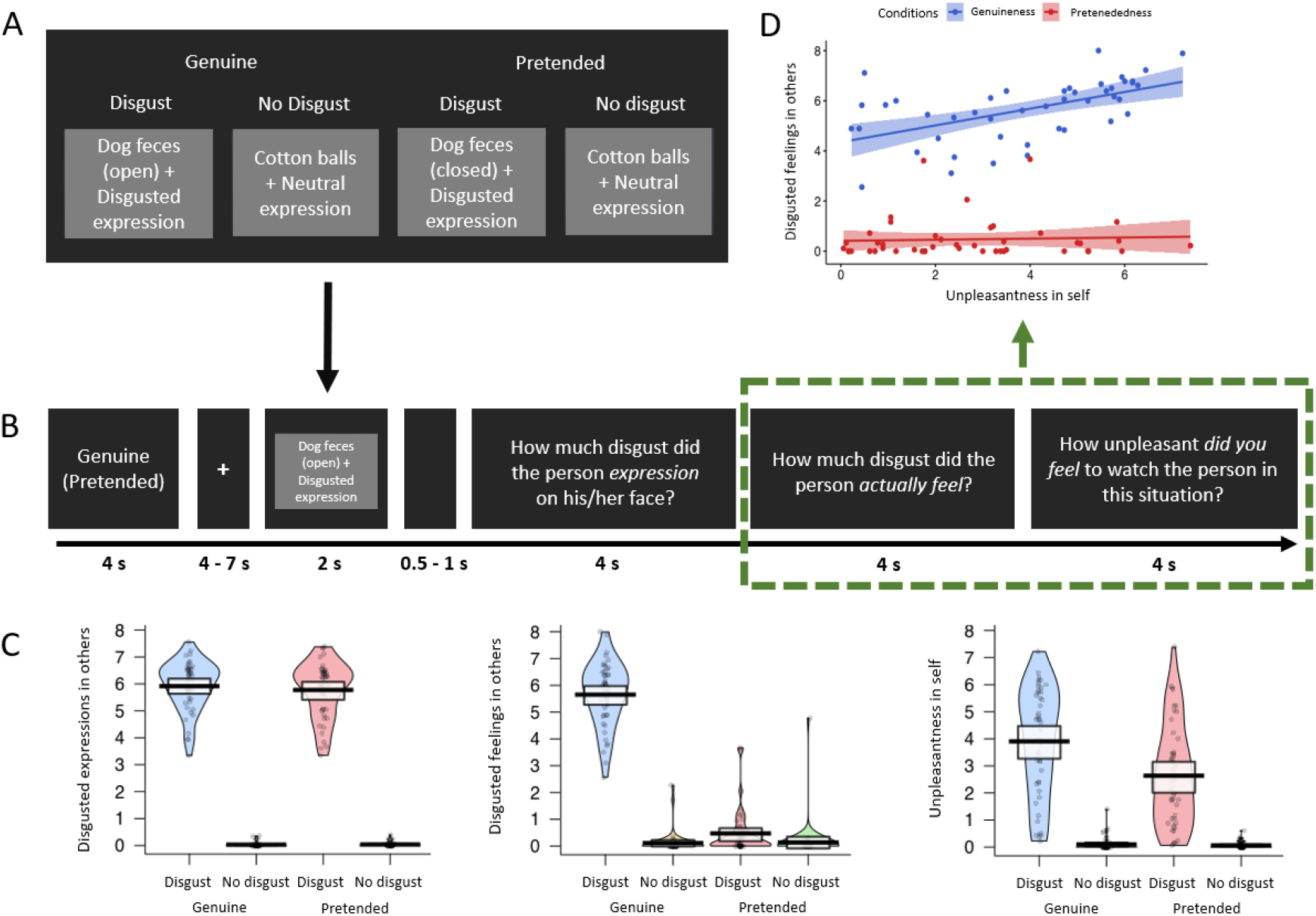
fMRI experimental design and behavioral results. (A) Overview of the experimental design with the four conditions genuine vs. pretended, disgust vs. no disgust. Examples show static images, while in the experiment, participants were shown video clips. (B) Overview of experimental timeline. At the outset of each block, a reminder of “genuine” or “pretended” was shown (both terms are shown here for illustrative purposes, in the experiment either genuine or pretended was displayed). After a fixation cross, a video in the corresponding condition appeared on the screen. Followed by a short jitter, three questions about the video were separately presented and had to be rated on a visual analog scale. These would then be followed by the next video clip and questions (not shown). (C) Violin plots of the three types of ratings for all conditions. No difference was found for the rating of disgusted expressions in others between the genuine disgust condition and the pretended disgust condition. For the ratings of disgusted feelings in others and unpleasantness in self, participants demonstrated higher ratings for genuine disgust than pretended disgust. Ratings of all three questions were higher in the disgusted situation than in the neutral situation, regardless of whether in the genuine or pretended condition. The thick black lines illustrate mean values, and the white boxes indicate a 95% CI. The dots are individual data, and the “violin” outlines illustrate their estimated density at different points of the scale. (D) Correlations of disgusted feelings in others and unpleasantness in self for the genuine disgust and the pretended disgust (the relevant questions were highlighted with a green rectangle). Results revealed a significant Pearson correlation between the two questions for the genuine disgust condition, but no correlation in the pretended disgust condition. The lines represent the fitted regression lines, bands indicate a 95% CI.

After the scanner session, participants came on another day to complete three questionnaires in the lab: the Empathy Components Questionnaire (ECQ), the Interpersonal Reactivity Index (IRI), and the Toronto Alexithymia Scale (TAS). The ECQ is categorized into Five subscales with 27 items (i.e., cognitive ability, cognitive drive, affective ability, affective drive, and affective reactivity), using a 4- point Likert scale ranging from 1 (“strongly disagree”) to 4 (“strongly agree”) (Batchelder, 2015; Batchelder et al., 2017). The IRI is divided into four subscales with 28 items (i.e., perspective taking, fantasy, empathic concern, and personal distress), using a 5-point Likert scale ranging from 0 (“does not describe me well”) to 4 (“describes me very well”) (Davis, 1980). The TAS is composed of three subscales with 20 items (i.e., difficulty describing feelings, difficulty identifying feelings, and externally oriented thinking), using a 5-point Likert scale ranging from 1 (“strongly disagree”) to 5 (“strongly agree”) (Bagby et al., 1994). Participants were debriefed at the end of the whole study.

### Behavioral data analysis

Repeated-measures ANOVAs were run in SPSS (version 26.0; IBM) to investigate the main effects and the interaction of the two factors genuine vs. pretended and disgust vs. no disgust. Furthermore, we conducted Pearson correlations to examine whether ratings of disgust feelings in others were correlated with unpleasantness in self for the genuine disgust and the pretended disgust. The comparison of the correlation coefficients was performed using a bootstrap approach with the R package bootcorci (https://github.com/GRousselet/bootcorci).

### fMRI data acquisition

We used a Siemens Magnetom Skyra 3T MRI scanner (Siemens, Erlangen, Germany) with a 32- channel head coil to collect fMRI data. A multiband-accelerated T2*-weighted echoplanar imaging (EPI) sequence was applied to collect functional whole-brain scans (TR = 1200 ms, TE = 34 ms, acquisition matrix = 96 × 96 voxels, FOV = 210 × 210 mm^2^, flip angle = 66°, inter-slice gap = 0.4 mm, voxel size = 2.2 × 2.2 × 2 mm^3^, multiband acceleration factor = 4, interleaved ascending acquisition in multi-slice mode, 52 slices co-planar to the connecting line between anterior and posterior commissure). Each of the two functional imaging runs lasted around 16 min (∼800 images per run). A magnetization-prepared rapid gradient-echo (MPRAGE) sequence was implemented to acquire structural images (TE/TR = 2.43/2300 ms, FOV= 240 × 240 mm^2^, flip angle = 8°, voxel size = 0.8×0.8×0.8 mm^3^, slice thickness = 0.8 mm, ascending acquisition, 208 sagittal slices, single-shot multi-slice mode).

### fMRI data processing and mass-univariate functional segregation analyses

Imaging data preprocessing was performed with a combination of Nipype (Gorgolewski et al., 2011) and MATLAB (version R2018b 9.5.0; MathWorks) with Statistical Parametric Mapping (SPM12; http://www.fil.ion.ucl.ac.uk/spm/software/spm12/). Raw data were arranged into BIDS format (http://bids.neuroimaging.io/; Gorgolewski et al., 2016). Functional data were 1) slice time corrected to the middle slice (Sladky et al., 2011), 2) realigned to the first image of each session, 3) co-registered to the T1 image, 4) segmented between grey matter, white matter, and cerebrospinal fluid (CSF), 5) normalized to MNI template space using Diffeomorphic Anatomical Registration Through Exponentiated Lie Algebra (DARTEL) toolbox (Ashburner, 2007), and 6) smoothed using a 6 mm full width at half-maximum (FWHM) of the Gaussian kernel. To improve data quality, scrubbing was performed when the frame-wise displacement (FD) of a scan was larger than 0.5 mm (Power et al., 2012; Power et al., 2014).

In order to perform mass-univariate functional segregation analyses, we created a first-level GLM design matrix composed of two identically modeled runs for each participant. Seven regressors of interest were entered in each model: stimulation phase of the four conditions (i.e., genuine disgust, genuine no disgust, pretended disgust, pretended no disgust; 2000 ms), rating phase of the three questions (i.e., disgusted expressions in others, disgusted feelings in others, and unpleasantness in self; 12000 ms). Six head motion parameters and the scrubbing regressors (FD > 0.5 mm; if applicable) were additionally entered as nuisance regressors.

On the second level, we used a flexible factorial design for the group-level analysis. Three factors were included: a between-subject factor (i.e., subject) that was specified independent and with equal variance, a within-subject factor (i.e., genuine or pretended) that was specified dependent and with equal variance, and a second within-subject factor (i.e., disgust or no disgust) that was specified dependent and with equal variance were included in the design (Gläscher & Gitelman, 2008). Four contrasts were computed: 1) genuine: disgust – no disgust, 2) pretended: disgust – no disgust, 3) genuine disgust – pretended disgust, and 4) genuine (disgust – no disgust) – pretended (disgust – no disgust).

An initial threshold of *p*< 0.001 (uncorrected) at the voxel level and a family-wise error (FWE) correction (*p* <⍰0.05) at the cluster level were applied. The cluster extent threshold was determined by the SPM extension “cp_cluster_Pthresh.m” (https://goo.gl/kjVydz).

### Brain-behavior relationships

We built a multiple regression model on the group level to investigate the relationship between specific brain activations and behavioral ratings. In this model, the contrast genuine disgust – pretended disgust was set as the dependent variable, and differences between conditions for three behavioral ratings were specified as independent variables. The reason that we entered the condition differences for both brain signals and behavioral ratings into the analyses was to control for the potential effects of perceptual salience. Moreover, we used the contrast genuine disgust – pretended disgust instead of the more exhaustive contrast genuine (disgust – no disgust) – pretended (disgust – no disgust), because our aim was to focus on the genuine and pretended disgust conditions rather than the neutral conditions. In addition, the disgust contrast showed more robust (in terms of statistical effect size) and widespread activations across the brain, making it more likely to pick up possible brain-behavior relationships. The same threshold as above (i.e., cluster-wise FWE correction, *p* <⍰0.05) was applied in this analysis. All covariates were mean-centered. An intercept was added to the model.

### Analyses using dynamic causal modeling (DCM)

We considered the following regions of interest (ROI) for the DCM model space: the right aIns and rSMG according to the previous study of pain (Zhao et al., 2021b) and the left (primary) olfactory cortex according to the exploratory analyses. As for the latter, the results showed no evidence that effective connectivity between aIns and rSMG for genuine disgust vs. pretended disgust was distinct. Therefore, we extended the analysis to the primary olfactory cortex, which was not hypothesized when planning this study but highly plausible given the employed task and the specific link between olfaction and disgust. In fact, previous studies indeed demonstrated the olfactory cortex was not only engaged in perceptual processes (e.g., odor perception and recognition), but also in affective processing of disgust-related experiences (Gottfried et al., 2002; Zelano et al., 2011; Alessandrini et al., 2016; Schulze et al., 2017; Schienle et al., 2020). We additionally defined an ROI of the right olfactory cortex as a comparison to the left olfactory cortex. The ROI masks were defined as the anatomical masks created by the Wake Forest University (WFU) Pick Atlas SPM toolbox (http://fmri.wfubmc.edu) with the automated anatomical atlas (AAL). Note that the olfactory cortex mask defined in AAL largely overlaps with the primary olfactory cortex that we were interested in (Desikan et al., 2006).

Three DCM analyses were performed based on different considerations. Firstly, to investigate if the distinct effective connectivity between aIns and rSMG we have found between genuine pain and pretend pain could be observed in disgust as well, we performed a DCM analysis between the right aIns and rSMG under the manipulation of genuine disgust and pretended disgust. Secondly, to explore if other brain patterns could dissociate genuine disgust and pretended disgust, we performed a second DCM analysis between the right aIns and the left olfactory cortex. Finally, to test the robustness of the second DCM model, we performed a third DCM analysis between the right aIns and the right olfactory cortex.

All DCM analyses were performed with DCM12.5 implemented in SPM12 (v. 7771). As a first step, individual time series were extracted separately for each ROI. The voxels were determined both on a group-level and an individual-level threshold to ensure the selected voxel were indeed engaged in a task-relevant activity instead of random signal fluctuations (Holmes et al., 2020). The initial threshold was set as *p* < 0.05, uncorrected. The significant voxels in the main effect of genuine disgust and pretended disgust were further selected by an individual threshold. An individual peak coordinate within the ROI mask was searched for each participant and an individual mask was consequently defined using a sphere of the 6 mm radius around the peak. The individual time series for each ROI was subsequently extracted from the significant voxels of the individual mask and summarized by the first eigenvariate. For the second and third DCM analyses, seven participants were excluded as no voxels survived significance testing in either the left or right olfactory cortex. In the next step, three regressors of interest were specified: genuine disgust, pretended disgust, and the video input condition (the combination of genuine disgust and pretended disgust). The reasons for not specifying the no-disgust conditions were that 1) disgust conditions were our main focus, and 2) adding effects of non-interest would inevitably increase the model complexity. In DCM, three sets of parameters were estimated: bidirectional connections between the regions and their self- connections (matrix A), modulatory effects (i.e., genuine disgust and pretended disgust) on the between-region connections (matrix B), and driving inputs (i.e., the video input condition) on both regions (matrix C) (Zeidman et al., 2019a). Finally, we performed group-level DCM inference using parametric empirical Bayes (Zeidman et al., 2019b). An automatic search was conducted over the entire model space (max. n =256) using Bayesian model reduction (BMR) and random-effects Bayesian model averaging (BMA), resulting in a final group model that takes accuracy, complexity, and uncertainty into account (Zeidman et al., 2019b). This procedure was similarly performed for all three DCM analyses. We reported all parameters with positive evidence on the posterior probability (*pp* > 0.75). Finally, modulatory effects of the genuine and pretended disgust conditions were compared using a paired sample t-test for each group-averaged model.

To probe whether task-related modulatory effects were associated with behavioral measurements, we performed multiple linear regression analyses of modulatory parameters with, 1) the three behavioral ratings, and 2) the empathy-related questionnaires (i.e., IRI, ECQ, and TAS). We set up two regression models for the genuine and pretended disgust conditions, respectively, in which the DCM parameters of modulatory effects were determined as dependent variables and the three ratings as independent variables. Considering that interactions between behavioral ratings might contribute to the regression model, five regression models (with and without interaction) were tested for both conditions. Results (Supplementary Table 1) showed the model without any interaction outperformed the other models for both genuine disgust and pretended disgust. We thus reported the results of the winning multiple regression model in the results section. We performed two additional regression models for both conditions in which DCM modulatory effects were set as dependent variables and scores of each questionnaire subscale were set as independent variables, respectively. Given the number of independent variables was considerable (>10), we used a stepwise regression approach to perform the analyses for questionnaires. As two participants did not complete all three questionnaires, we excluded their data from the regression analyses. The statistical significance of the regression analysis was set to *p* < 0.05. The multicollinearity for independent variables was diagnosed using the variance inflation factor (VIF) that measures the correlation among independent variables, in the R package car (https://cran.r-project.org/web/packages/car/index.html). Here a rather conservative threshold of *VIF* < 5 was adapted as an indication of no severe multicollinearity (Menard, 2002; James et al., 2013).

## Results

### Behavioral results

We performed three repeated-measures ANOVAs with the factors *genuineness* (genuine vs. pretended) and *disgust* (disgust vs. no disgust), for each of the three behavioral ratings. For ratings of disgusted *expressions* in others (Figure 1C, left), the main effect of the factor genuineness was not significant: *F* _genuineness_ (1, 44) = 1.861, *p* = 0.179, η^2^ = 0.041. There was a main effect of disgust: participants showed higher ratings for the disgust vs. no disgust conditions, *F* _disgust_ (1, 44) = 1769.396, *p* < 0.001, η^2^ = 0.976. The interaction term was not significant, *F* _interaction_ (1, 44) = 2.270, *p* = 0.139, η^2^ = 0.049. For ratings of disgusted feelings in others (Figure 1C, middle), there was a main effect of genuineness: participants showed higher ratings for the genuine vs. pretended conditions, *F* _genuineness_ (1, 44) = 510.686, *p* < 0.001, η^2^ = 0.921. There was also a main effect of disgust, as participants showed higher ratings for the disgust vs. no disgust conditions, *F* _disgust_ (1, 44) = 854.136, *p* < 0.001, η^2^ = 0.951. The interaction was significant as well, *F* _interaction_ (1, 44) = 360.516, *p* < 0.001, η^2^ = 0.891, and this was related to higher ratings of disgusted feelings in others for the genuine disgust compared to the pretended disgust condition. For ratings of unpleasantness in self (Figure 1C, right), there was a main effect of genuineness: participants showed higher ratings for the genuine vs. pretended conditions, *F* _genuineness_ (1, 44) = 37.694, *p* < 0.001, η^2^ = 0.461. There was also a main effect of disgust: participants showed higher ratings for the disgust vs. no disgust conditions, *F* _disgust_ (1, 44) = 141.277, *p* < 0.001, η^2^ = 0.763. The interaction was significant as well, *F* _interaction_ (1, 44) = 32.341, *p* < 0.001, η^2^ = 0.424, and this was related to higher ratings of unpleasantness in self for the genuine disgust compared to the pretended disgust condition. In sum, the behavioral data indicated that there was no difference in ratings of disgusted expression in others between the genuine and pretended disgust conditions, while higher ratings and large effect sizes of disgusted feelings in others and unpleasantness in self for the genuine disgust condition as compared to the pretended disgust condition. Moreover, results showed that participants rated expressions of disgust as more unpleasant than neutral expressions, regardless of whether they were genuine or not. These results were perfectly in line with our hypotheses and what we found in the pilot study.

We also found significant correlations between behavioral ratings of disgusted feelings in others and unpleasantness in self for the genuine disgust condition, *r* = 0.548, *p* < 0.001; while for the pretended disgust condition, the correlation was not significant, *r* = 0.051, *p* = 0.740 (Figure 1D). A bootstrapping comparison showed a significant difference between the two correlation coefficients, *p* = 0.025, 95% Confidence Interval (CI) = [0.073, 0.860].

### fMRI results: mass-univariate analysis

We computed four contrasts: 1) genuine: disgust – no disgust, 2) pretended: disgust – no disgust, 3) genuine disgust – pretended disgust, and 4) genuine (disgust – no disgust) – pretended (disgust – no disgust). In the first two contrasts, we found the predicted activations in bilateral aIns, aMCC, and rSMG, as well as significant (not originally predicted) activation in the olfactory cortex; in the third contrast, we found significant activation in the right aIns, as well as strong activation (k = 255) in the left olfactory cortex; in the last contrast, the only significant activation was found in the right cerebellum (Figure 2A and Table 1).

**Figure 2.**
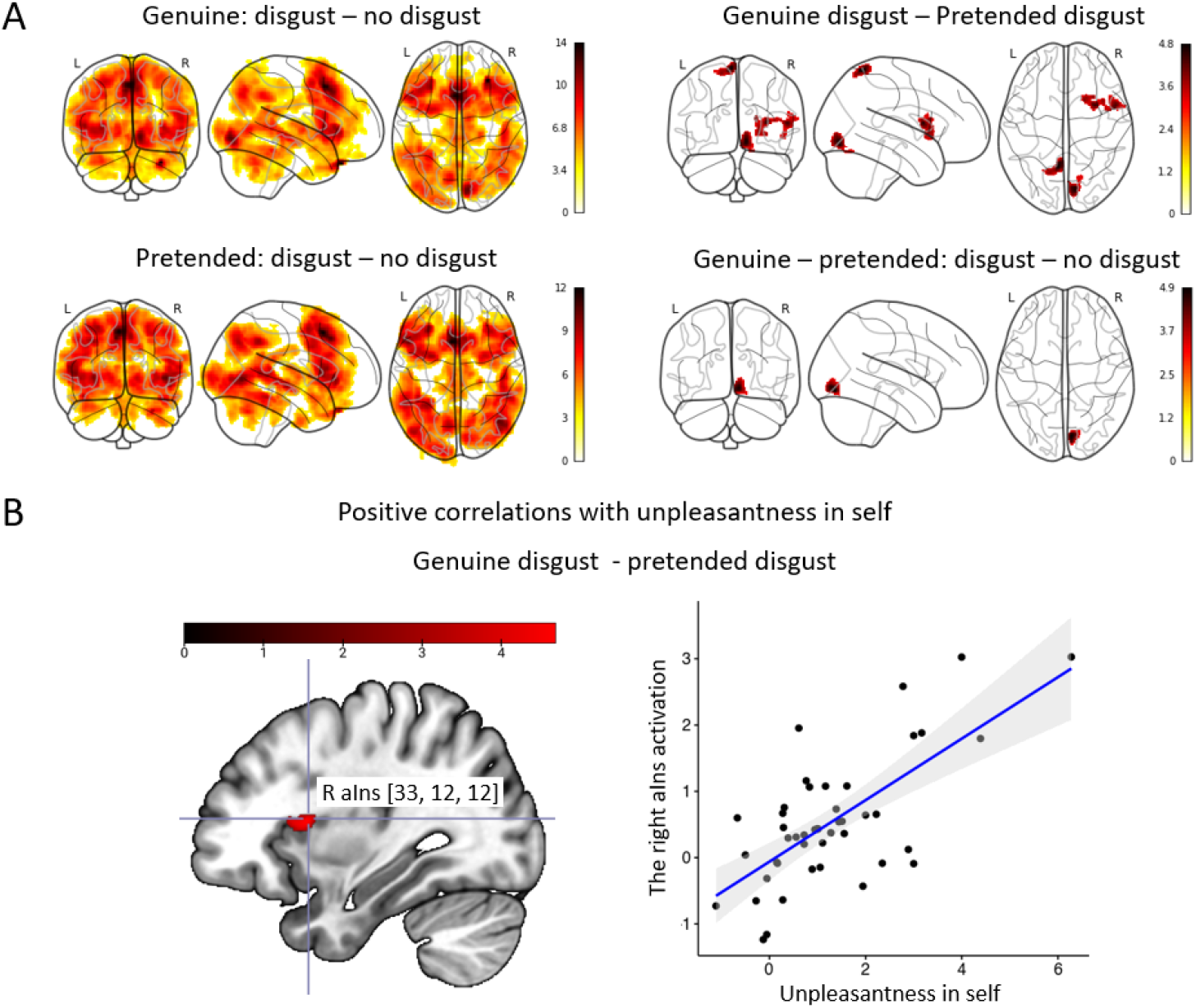
Neuroimaging results: Mass-univariate analyses. (A) Activation maps of genuine: disgust – no disgust (top left), pretended: disgust - no disgust (bottom left), genuine disgust – pretended disgust (top right), and genuine (disgust – no disgust) – pretended (disgust – no disgust) (bottom right). For contrasts of disgust –no disgust in both genuine and pretended conditions, we found expected brain activations in bilateral aIns, aMCC, and rSMG, and significant activation in the olfactory cortex; for the contrast of genuine disgust vs. pretended disgust, we found significant activation in the right aIns and strong activation in the left olfactory cortex (a cluster of k=255, though not pass the threshold); for the contrast of genuine (disgust – no disgust) vs. pretended (disgust – no disgust), the only significant activation was in the right cerebellum. (B) The multiple regression analysis demonstrated a significant cluster in the right aIns (peak: [33, 12, 12]) that was positively associated with the ratings of unpleasantness in self but not associated with the ratings of either disgusted expressions in others or disgusted feelings in others when comparing genuine disgust vs. pretended disgust. All activations are thresholded with cluster-level FWE correction, *p* < 0.05 (*p* < 0.001 uncorrected initial selection threshold). The lines of the scatterplots represent the fitted regression lines, bands indicate a 95% CI.

**Table 1.** Results of mass-univariate functional segregation analyses in the MNI space. Region names were labeled with the AAL atlas and thresholded with cluster-wise FWE correction, *p* < 0.05 (initial selection threshold *p* < 0.001, uncorrected). BA = Brodmann area, L = left hemisphere, R = right hemisphere.

| Region label | BA | Cluster size | x | y | z | t-value |
| --- | --- | --- | --- | --- | --- | --- |
| **1) Genuine: disgust - no disgust** |  |  |  |  |  |  |
| Temporal_Pole_Sup_R | 38 | 164988 | 32 | 33 | -33 | 13.57 |
| Supp_Motor_Area_L | 8 |  | -3 | 16 | 50 | 13.31 |
| Lingual_R | 18 |  | 9 | -84 | -6 | 11.89 |
| Frontal_Sup_Medial_R | 8 |  | 6 | 21 | 45 | 10.95 |
| Insula_L | 45 |  | -32 | 27 | 4 | 10.81 |
| Frontal_Inf_Oper_L | 44 |  | -50 | 15 | 6 | 10.80 |
| Insula_R | 13 |  | 33 | 27 | 4 | 10.17 |
| Frontal_Inf_Tri_L | 45 |  | -30 | 32 | 0 | 10.12 |
| Frontal_Inf_Oper_R | 44 |  | 52 | 15 | 15 | 9.80 |
| Frontal_Inf_Orb_R | 47 |  | 32 | 32 | -3 | 9.69 |
| Lingual_L | 17 | 422 | -21 | -66 | 4 | 5.10 |
| **2) Pretended: disgust - no disgust** |  |  |  |  |  |  |
| Supp_Motor_Area_L | 8 | 137060 | -4 | 16 | 50 | 11.95 |
| Frontal_Sup_Medial_L | 8 |  | -8 | 26 | 44 | 10.69 |
| Temporal_Mid_R | 19 |  | 44 | -68 | 2 | 10.35 |
| Frontal_Inf_Oper_L | 44 |  | -51 | 16 | 6 | 10.01 |
| Insula_L | 45 |  | -30 | 30 | 2 | 9.77 |
| Frontal_Inf_Tri_L | 45 |  | -56 | 20 | 12 | 9.68 |
| Cingulum_Mid_R | 8 |  | 8 | 20 | 45 | 9.47 |
| Parietal_Inf_L | 39 |  | -32 | -51 | 40 | 9.37 |
| Temporal_Pole_Sup_R | 38 |  | 32 | 34 | -33 | 9.35 |
| Frontal_Mid_L | 6 |  | -27 | 2 | 54 | 9.26 |
| Cingulate_Post_L | 23 | 1821 | -4 | -42 | 22 | 5.71 |
| Cingulate_Mid_R | 23 |  | -3 | -26 | 27 | 5.58 |
| Cingulate_Post_R | 23 |  | 8 | -39 | 22 | 5.33 |
| Cingulate_Mid_L | 24 |  | -3 | -12 | 30 | 5.23 |
| Vermis_9 | 37 | 522 | 2 | -57 | -39 | 5.27 |
| Cerebelum_9_L | 18 |  | -2 | -60 | -46 | 4.55 |
| Cerebelum_9_R | 37 |  | 10 | -57 | -51 | 3.96 |
| Temporal_Inf_L | 20 | 517 | -40 | -9 | -38 | 4.80 |
| Temporal_Pole_Mid_L | 38 |  | -28 | 8 | -39 | 3.62 |
| Cerebelum_Crus2_L | 18 | 487 | -8 | -76 | -34 | 5.15 |
| Cerebelum_Crus1_L | 18 |  | -18 | -80 | -27 | 3.21 |
| **3) Genuine disgust – pretended disgust** |  |  |  |  |  |  |
| Insula_R | 44 | 976 | 32 | 8 | 12 | 4.56 |
| Rolandic_Oper_R | 44 |  | 54 | 8 | 10 | 4.38 |
| Caudate_R | 48 |  | 21 | 14 | 15 | 4.05 |
| Putamen_R | 49 |  | 30 | 16 | -2 | 3.89 |
| Frontal_Inf_Oper_R | 44 |  | 40 | 12 | 14 | 3.85 |
| Lingual_R | 18 | 550 | 10 | -81 | -9 | 4.72 |
| Cerebelum_6_R | 18 |  | 15 | -70 | -18 | 3.35 |
| Precuneus_L | 7 | 474 | -6 | -56 | 69 | 4.85 |
| Parietal_Sup_L | 7 |  | -16 | -63 | 64 | 3.95 |
| **4) Genuine (disgust – no disgust) – pretended (disgust – no disgust)** |  |  |  |  |  |  |
| Lingual_R | 18 | 431 | 8 | -81 | -10 | 4.90 |

To identify whether or which brain activity was selectively related to the behavioral ratings described above, we performed a multiple regression analysis on the whole brain level where we explored the relationship of activation in the contrast genuine disgust – pretended disgust with the three behavioral ratings. The only significant cluster we found encompassed the right aIns, extending into the right inferior frontal gyrus, and this was selectively related to ratings of self-unpleasantness (Figure 2B) rather than the ratings of disgusted expressions in others or the disgusted feelings in others.

### DCM results

We first performed a DCM analysis of the effective connectivity between the right aIns and rSMG to examine if the group-averaged model replicated what we found in our previous study on pain (Zhao et al., 2021b). Specifically, we focused on the modulatory effect of genuineness, namely, whether the experimental manipulation of genuine disgust vs. pretended disgust tuned the bidirectional neural dynamics from aIns to rSMG and *vice versa*, in terms of both directionality (sign of the DCM posterior parameter) and intensity (magnitude of the DCM posterior parameter). If the experimental manipulation modulated the effective connectivity, we would observe a positive posterior probability (*p*_*p*_ > 0.75) of the modulatory effect. The reasons that we did not include aMCC in this analysis were that 1) unlike aIns, in aMCC we did not find strong evidence for the task involvement (in the univariate and multiple regression analyses), 2) for comparability of the present model with the previous model on pain, where aMCC had not been included either.

Similar to what we found in the pain study, strong evidence (*pp* > 0.95; *pp* = 1.00) of inhibitory modulatory effects on the aIns-to-rSMG connection was shown for both the genuine disgust condition and the pretended disgust condition (see Figure 3A). However, we did not find a significant difference when comparing the strength of these two modulatory effects, *t*_*44*_ = -1.045, *p* = 0.302 (Mean _genuine disgust_ = -1.214, 95% CI = [-1.462, -0.927]; Mean _pretended disgust_ = -1.095, 95% CI = [-1.361, - 0.799]. Note that the mean values in the test could slightly differ from those shown in the DCM models of Figure 3, since we used frequentist statistics for comparison analysis rather than the Bayesian approach that was implemented to compute the parameters for the DCM model. We did not find robust evidence on the intrinsic connectivity either from aIns to rSMG or *vice versa*. Moreover, there was no evidence of a modulatory effect on the rSMG to aIns connection, which was in line with what we had found for pain. Taken together, the DCM analysis between aIns and rSMG partially replicated the results of the pain study, namely the inhibitory modulatory effect from aIns to rSMG for both genuine and pretended conditions; however, this inhibitory modulatory effect failed to dissociate the experimental manipulation of genuine disgust and pretended disgust, suggesting that a distinctive pattern or set of brain regions underpins how the genuineness of disgust is processed.

**Figure 3.**
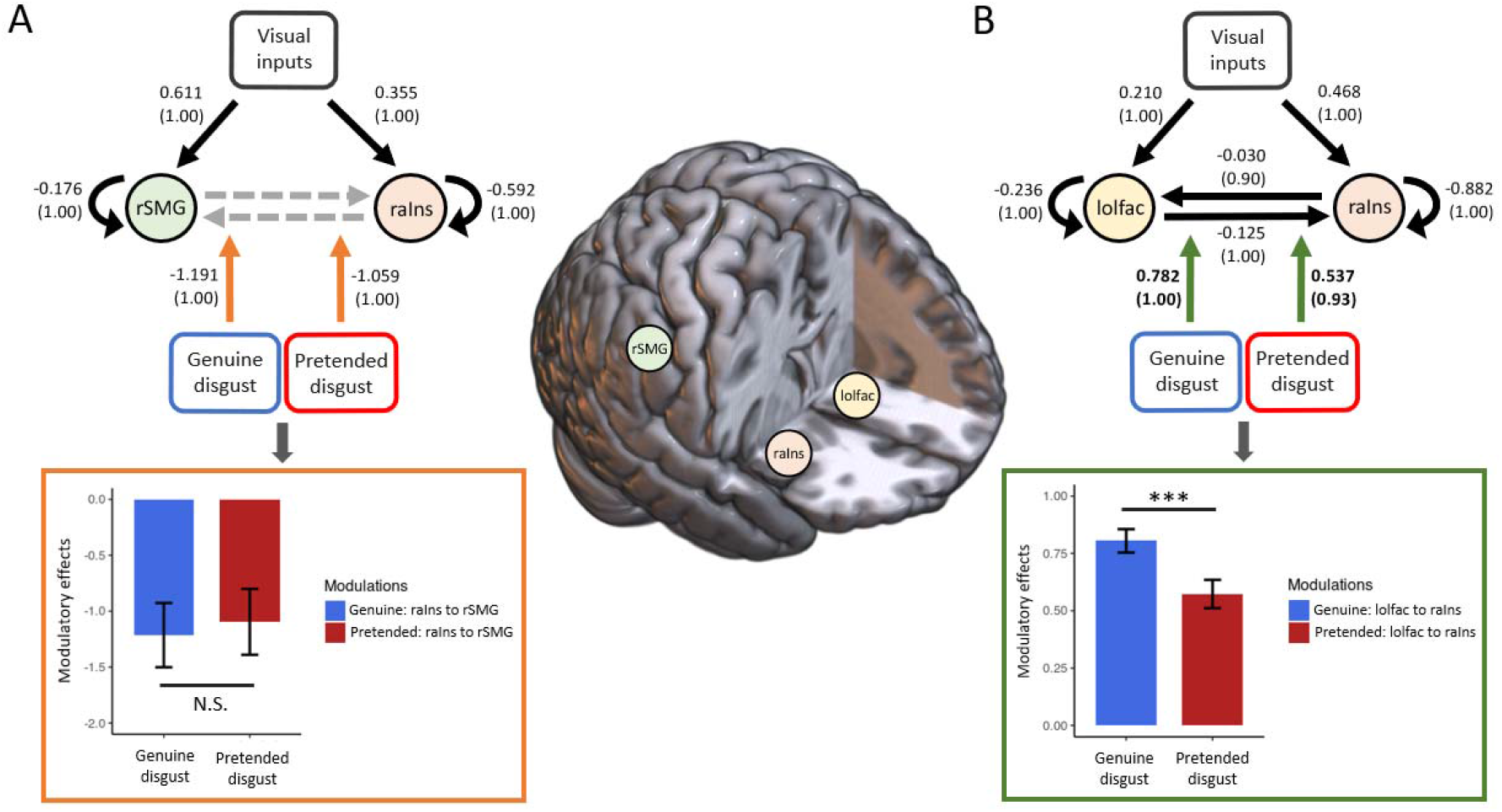
DCM results and brain-behavior analyses. Three-dimensional visualization of ROIs involved in the two DCM analyses is shown in the upper middle. (A) The group-average DCM model of the right anterior insula and the right supramarginal gyrus (rSMG) for genuine disgust and pretended disgust. We found inhibitory modulatory effects (orange arrows) for both conditions. All DCM parameters of the optimal model showed greater than a 99% posterior probability (very strong evidence) except the bi-directionally intrinsic connectivity between the right aIns (raIns) and rSMG (grey dashed arrow; no evidence of existence, *pp* < 0.50). Paired sample *t*-test showed no difference in the inhibitory modulatory effects on the raIns-to-rSMG connection between genuine disgust and pretended disgust. This result is highlighted with an orange rectangle. Data are mean ± 95% CI. (B) The group-average DCM model of the raIns and the left olfactory cortex (lolfac) for genuine disgust and pretended disgust. We found the excitatory modulatory effect (green arrows) for both conditions. All DCM parameters of the optimal model showed greater than or equal to a 90% posterior probability (*pp* > 0.75, positive evidence). Paired sample *t*-test showed a stronger excitatory modulatory effect of the lolfac-to-raIns connection for genuine disgust as compared to pretended disgust (*** *p* < 0.001). This result is highlighted with a green rectangle. Data are mean ± 95% CI. For the DCM models, values without the bracket quantify the strength of connections; positive values indicate neural excitation and negative values indicate neural excitation. Values in the parentheses indicate the posterior probability of connections.

We therefore performed an exploratory DCM analysis to test whether distinct modulatory effects could be found for genuine and pretend disgust on the connection between the right aIns and the left olfactory cortex (see Figure 3B). As mentioned in the methods sections, while the involvement of the left olfactory cortex was mainly exploratory and data-driven, it was also plausible on theoretical grounds. Results showed a significant excitatory effect for both genuine disgust (strong evidence, *pp* = 1.00) and pretended disgust (positive evidence, *pp* = 0.93) on the connection of the left olfactory cortex to the right aIns. A further comparison analysis on the modulatory effect between conditions revealed a stronger excitatory modulatory effect for genuine disgust as opposed to pretended disgust, *t*_*37*_ = 4.450, *p* < 0.001 (Mean _genuine disgust_ = 0.805, 95% CI = [0.755, 0.851]; Mean _pretended disgust_ = 0.573, 95% CI = [0.517, 0.628]. We did not find any modulatory effect on the reverse connection of the right aIns to the left olfactory cortex.

Finally, to justify whether lateralization of the olfactory cortex largely influenced the robustness of the modulatory effect we found in the DCM model above, we performed an additional DCM analysis on the connection between the right aIns and the right olfactory cortex (see Supplementary Figure 1). Results showed a similar group-average model to that of the right aIns and the left olfactory cortex. Importantly, we replicated the excitatory modulatory effect on the connection of the olfactory cortex to aIns in the sense of significant evidence for both conditions (genuine disgust: positive evidence, *pp* = 0.91; pretended disgust: positive evidence, *pp* = 0.88). No evidence of the modulatory effect on the connection of aIns to the right olfactory cortex was found, which was consistent with the group-average model with the left olfactory cortex. A further comparison analysis did not find significant difference on the strength of the two modulatory effects, *t*_*37*_ = 0.595, *p* = 0.556 (Mean _genuine disgust_ = 0.624, 95% CI = [0.503, 0.738]; Mean _pretended disgust_ = 0.577, 95% CI = [0.488, 0.678]).

### Individual associations between modulatory effects, behavioral ratings, and questionnaires

Two linear regression models were computed to examine how the excitatory modulatory effect on the connection of the (left) olfactory cortex to aIns was related to behavioral ratings respectively for genuine disgust and pretended disgust. Results showed that none of the ratings was significant for either the genuine disgust model or the pretended disgust model. No severe collinearity problem was detected for either regression model (all *VIFs* < 4.500; the smallest *VIF* =1.229 and the largest *VIF* = 4.410).

Another two linear regression models were tested to investigate whether subscales of all three questionnaires could explain the excitatory modulatory effect for genuine disgust and pretended disgust. For the genuine disgust condition, we found that the modulatory effect was significantly explained by scores of the perspective-taking subscale of the IRI: *F*_model_ (1, 35) = 4.177, *p* = 0.049, *R*^*2*^ = 0.109; *B* = 0.011, *beta* = 0.331, *p* = 0.049. No significant predictor was found with any subscale in the other two questionnaires (i.e., ECQ and TAS). None of the three questionnaires significantly explained variations of the modulatory effect for the pretended disgust condition. No severe collinearity problem was detected for either regression model (all *VIFs* < 2.500; the smallest *VIF* =1.000 and the largest *VIF* = 2.198).

## Discussion

Using a paradigm matched to our previously published study on pain (Zhao et al., 2021b), and in the same sample and experimental session, we here report how participants responded to video clips presenting people who supposedly either genuinely experienced disgust or merely pretended to feel disgusted. Combining mass-univariate analysis with effective connectivity (DCM) analyses, we aimed to clarify two main questions: 1) whether neural responses in areas such as aIns and aMCC to the disgust of others were indeed related to a veridical sharing of affect, as opposed to simply tracking sensory-driven responses to salient affective displays, and 2) whether the effective connectivity between aIns and rSMG that we previously found to disentangle genuine pain from pretended pain also enabled the dissociation of genuine disgust vs. pretended disgust.

We found increased activations in the right aIns for genuine disgust as compared to pretended disgust that was selectively associated with the unpleasantness in self. These findings are in line with what we have found in pain (Zhao et al., 2021b), implying an essential role of aIns in the processing of shared feelings with others for both pain and disgust. However, an intriguing question is if the aIns activation we observed in these two aversive states reflects a form of cross-modal affective processing, or rather modality-dependent affective experiences? A study using multi-voxel pattern analysis (MVPA) has shown both cross-modal and modality-specific evidence in terms of the subfields of aIns for pain and disgust: in the left aIns (and aMCC), the shared encoding was detected for first-hand and vicarious pain and disgust, regardless of the same or different modality; while in the right aIns, sensory-specific rather than modality-independent patterns were more plausible for processing first-hand and vicarious pain and disgust (Corradi-Dell’Acqua et al., 2016). Taken together, the aIns activation we found suggests the engagement of affective processing that was related to others’ pain and disgust, while future research that explicitly matches pain and disgust salience is required to further investigate whether this activation indicates cross-modal or modality-dependent affective experiences.

We found significant inhibitory modulatory effects on the connection of aIns to rSMG for both genuine and pretended disgust, but these effects did not differ significantly. This implies that we only partially replicate the findings of the pain study: while we reproduce a role of the aIns and rSMG connectivity, their crosstalk does not explain the distinction between genuine and pretended expressions of disgust, as is the case for pain (Zhao et al., 2021b). We speculate that the absence of differences between two conditions in rSMG activation as well as the inhibitory modulatory effect could be related to generally lower salience of aversive experiences in the disgust task compared to that of pain. Further investigation is required to test this assumption.

Contrary to our expectations, we did not find any significant activation in rSMG for genuine disgust as compared to pretended disgust; instead, we showed a relatively stronger engagement of the primary olfactory cortex between conditions. As we mentioned beforehand, we did not target the latter area when planning the study but later included it inspired by the exploratory analysis. The (primary) olfactory cortex has been considered to mainly comprise the anterior olfactory nucleus, the olfactory tubercle, piriform cortices, and subregions of amygdala and entorhinal cortex (Savic et al., 2000; Tzourio-Mazoyer et al., 2002; G. Zhou et al., 2019). Studies have found that this region is recruited not only for direct olfactory sensations but also for the indirect experience of olfactory processing, such as odor imagery (Djordjevic et al., 2005; Bensafi et al., 2007) and odor prediction (Zelano et al., 2011). Olfactory priming could facilitate the identification of the emotion of disgust (Seubert et al., 2010a; Seubert et al., 2010b); in turn, priming with a disgusted face compared to a happy face enhanced activation in the olfactory cortex when processing pleasant odors (Schulze et al., 2017). These findings imply the engagement of the olfactory cortex in integrating olfactory processes with visually conveyed affective information. Furthermore, the primary olfactory cortex has been suggested to participate in processing the emotion of disgust, without necessarily experiencing sensory-related disgust. Compared with healthy controls, patients with reduced olfactory function (e.g., anosmia and hyposmia) have been found to identify less disgust for facial expressions of disgust and show greater activations in the primary olfactory cortex, suggestive of a compensatory effect, for disgusting scenes (Schienle et al., 2020). Altogether, the stronger engagement of the olfactory cortex might be related to a higher level of identified disgust in others when individuals observed others genuinely experiencing disgust compared with pretending disgusted.

The exploratory DCM analysis of the right aIns and the left olfactory cortex demonstrated a stronger excitatory modulatory effect on the olfactory cortex to aIns connection for genuine disgust as opposed to pretended disgust. Studies from nonhuman primates and humans using tractography have shown structural connections (for human: functional connectivity as well, see Deen et al., 2010) between the olfactory cortex and a partial region of aIns (Mufson & Mesulam, 1982; Carmichael et al., 1994; Ghaziri et al., 2015; Ghaziri et al., 2018). Specifically, as an important part of the secondary olfactory cortex, aIns is considered to engage in receiving and integrating the primary olfactory- affective information conveyed by the primary olfactory cortex. Moreover, the external sensory and affective messages seem to be already preprocessed in the primary olfactory cortex before they are conveyed to the secondary cortices (Soudry et al., 2011, for review; Seubert et al., 2013, for meta-analyses). Together with the evidence of higher brain activation in the right aIns and left olfactory cortex for genuine disgust compared to pretended disgust, we speculate that the increased excitatory modulatory effect on the olfactory-to-aIns connection may be related to the processing of passing messages of higher disgust emotion identified in others to activations related to affect processing, which may constitute the neural underpinning of the increased shared unpleasantness with others. This idea would also be in line with the theoretical framework proposed by Coll et al. (2017), that higher identified emotion in others contributes to stronger shared affect, and that the fully-fledged empathic response may be an integrated consequence of (at least) these two processes.

We performed another DCM analysis between the right aIns and the right olfactory cortex to test whether lateralization of the olfactory cortex had a large impact on the reliability of the modulatory effect we detected in DCM model with the left olfactory cortex. Results showed a very similar pattern to the DCM model with the left olfactory cortex, in the sense of replicating the excitatory modulatory effect on the connection of the olfactory cortex to aIns for both conditions and absence of any condition-dependent modulatory effect on the connection of the opposite direction. Even though for this model we did not find a significant difference in the modulatory effects between genuine disgust and pretended disgust, these results at least attest to the robustness of the modulatory effect from the olfactory cortex, regardless of the left or right hemisphere, to the right aIns.

We found the excitatory modulatory effect for genuine disgust was positively related to individual perspective-taking scores. This finding demonstrated that the connection of the olfactory cortex to aIns for genuine disgust was related to the tendency of adopting the psychological point of view of others, which would finally contribute to the level of one’s own affective responses to the emotion felt by another person. This view is supported by the evidence that aIns, especially the right aIns, is implicated in distinct neural patterns of representing self- and other-related aversive states for disgust as well as pain (Corradi-Dell’Acqua et al., 2016). No association with any questionnaire was found for pretended disgust. Taking all these results together, we would speculate that for the genuine disgust condition, the olfactory cortex interacts with aIns to achieve genuine identification of disgust in others. This would call for a higher demand to take the other’s perspective, and in this way may contribute to the higher shared affect. For the pretended pain condition, sensory-driven (“automatic”) emotion processing induced by the saliency of disgust expression interacts with the cognitive processes (i.e., knowing this person was merely acting out and did not feel any disgust at all), resulting in both a low level of identified disgust and shared affect. In this case, it may be less important to recruit the function of perspective-taking to share the emotions of others. However, further investigation is required to test these interpretations.

Note that, the aim of the current study, in relation to our previous work (Zhao et al., 2021b), was to investigate similar but parallel research questions in two closely related yet distinct modalities (i.e., disgust and pain). In this, we abstained from a quantitative comparison within an analysis framework that included data from both tasks for the following reasons. First, direct comparison would have required the saliency of the stimuli in the two modalities to be identical. Yet in an exploratory analysis, we found that the genuine pain always evoked higher ratings on all three behavioral measurements and stronger activations in regions that we were interested in, i.e., aIns, aMCC, and rSMG, as opposed to genuine disgust (results of this exploratory analysis are not reported in detail here; public links to the behavioral and functional data for disgust and pain can be found in the data availability statement and Zhao et al., 2021b). In this sense, any conclusion or inference drawn from directly comparing these two modalities could arguably be attributed to the difference in stimuli salience or the responses to that salience, rather than the essential emotional properties in pain and disgust. Second, incorporating all data into one large analysis model would have resulted in a very complex analysis approach, especially when considering the many different factors of our experimental design, the regions we were interested in, and the behavioral data we wanted to incorporate. While the rationale to analyze the tasks separately thus resulted in more robust evidence regarding each task, one limitation of the current study is that comparison across and replication across tasks allows insights into similarity and differences in principles and brain function rather in a descriptive way. Future research should thus better match the saliency (Sharvit et al., 2015), on both perceptual and affective levels, between pain and disgust to allow direct quantitative comparisons across modalities, while also keeping in mind the complexity of the experimental design.

In conclusion, the current study largely replicates, as well as expands our previously reported findings on pain. Firstly, and similar to what we have shown for empathy for pain using the same experimental approach and within the same study, we provide evidence that responses related to empathy for disgust in aIns can indeed be linked to the affective sharing rather than merely perceptual saliency. Secondly, we show how aIns and the olfactory cortex, instead of aIns and rSMG that we previously found in pain, orchestrate the tracking of disgust felt by another person. Taken together, these findings indicate that similar as well as distinct brain networks are engaged in processing different affective experiences, in this case pain and disgust, experienced by others. This refines and expands our understanding of the neural bases of empathy, from a dynamic and multi- modal perspective.

## Acknowledgements

This work was supported by Chinese Scholarship Council (CSC) Grant (201604910515) and Vienna Doctoral School in Cognition, Behavior and Neuroscience (VDS CoBeNe) completion grant fellowship to Y.Z.; the Vienna Science and Technology Fund (WWTF VRG13-007) to C.L., and the Austrian Science Fund (FWF P 32686) to C.L. and M.R..

## Notes

**Conflicts of interest:** The authors declare no competing financial interests.

### Competing Interest Statement

The authors have declared no competing interest.

