## Supplementary Figure 1 for "Effective connectivity reveals distinctive patterns in response to others’ genuine affective experience of disgust as compared to pain"

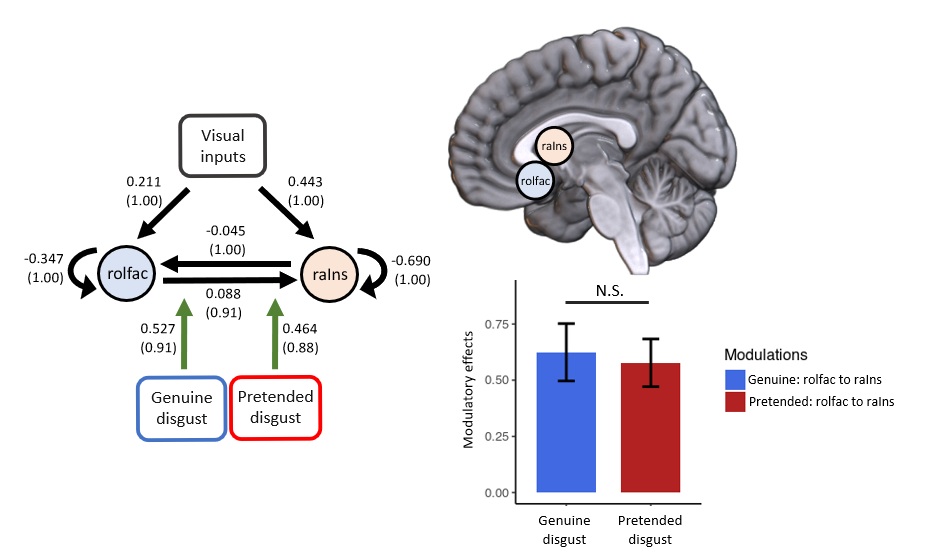

**Supplementary Figure 1.** The left panel: the group-average DCM model of the right anterior insula (raIns) and the right olfactory cortex (rolfac) for genuine disgust and pretended disgust. We found excitatory effects (green arrows) for both conditions. All DCM parameters of the optimal model showed greater than a 75% posterior probability (positive evidence). The top right panel: sagittal view of the two ROIs used for this DCM analysis. The bottom right panel: paired sample *t*-test showed no difference in the inhibitory effect on the rolfac-to-raIns connection between genuine disgust and pretended disgust. Data are mean ± 95% CI. For the DCM model, values without the bracket quantify the strength of connections; positive values indicate neural excitation and negative values indicate neural excitation. Values in the parentheses indicate the posterior probability of connections.
