## Supplementary Table 1 for "Effective connectivity reveals distinctive patterns in response to others’ genuine affective experience of disgust as compared to pain"

**Supplementary Table 1.** Model comparison of linear regression models with three behavioral ratings (independent variables) and the inhibitory effect (dependent variable) for genuine disgust and pretended disgust. Smaller AIC/BIC indicates better model fit. Results showed that M1 (without interaction; highlighted with underlining) was the best fitting model for both genuine disgust and pretended disgust.

| Regression Model | AIC | BIC |
| --- | --- | --- |
| Genuine disgust |  |  |
| M1 (expression + feeling + unpleasantness) | -27.422 | -19.234 |
| M2 (expression * feeling + unpleasantness) | -25.547 | -15.722 |
| M3 (expression + feeling * unpleasantness) | -25.515 | -15.690 |
| M4 (expression * unpleasantness + feeling) | -25.429 | -15.604 |
| M5 (expression * feeling * unpleasantness) | -21.604 | -6.865 |
| Pretended disgust |  |  |
| M1 (expression + feeling + unpleasantness) | -10.697 | -2.509 |
| M2 (expression * feeling + unpleasantness) | -10.650 | -0.824 |
| M3 (expression + feeling * unpleasantness) | -10.421 | -0.596 |
| M4 (expression * unpleasantness + feeling) | -9.5726 | 0.253 |
| M5 (expression * feeling * unpleasantness) | -5.4612 | 9.277 |
